# Person identity drives neural similarity more than action and valence during dynamic emotion perception

**DOI:** 10.64898/2026.06.01.728854

**Authors:** Deborah E. Okeke, Robert S. Chavez

## Abstract

Facial perception is a central feature of everyday social encounters and a rich source of emotional information. Classic functional magnetic resonance imaging (fMRI) studies of emotional facial processing used static photos emotional expression to identify regions of the brain showing univariate differences in response magnitudes between different emotional categories. However, there has been much less work identifying how the brain represents dynamic emotional facial expression and the factors that drive the similarity among these representations. In the current study, incorporated dynamic facial expression stimuli and representational similarity analysis to compare three competing hypothesized models of similarity of each of the stimuli presented: action being made, valence of the expression, and the identity of the person being perceived. Participants were shown short videos of fourteen volunteer actors making positive or negative facial expressions directed either toward or away from the camera. Activation patterns were compared against competing models on a trial-by-trial basis using a full multilevel modeling approach. Results showed that the identity of the person in the video was a greater predictor of brain responses similarity than the action or valence across widely distributed brain systems, particularly in the default mode network and lower-level visual processing regions. This suggests that the specific identity of the stimulus being perceived is a central driver of neural response similarity during perceptual encoding in dynamic facial processing.

---

Humans encode an endless array of co-occurring information. As a result, our brains have adapted to utilize complex systems to represent information that can be decoded and accessed in future encounters. These representational systems form the link between stimuli and the appropriately evoked response, both cognitive and physical. In a social context, these systems support our ability to simultaneously adapt to social attention demands, determine emotional valence and key person identifiers, among other processes. Often these neural representations overlap significantly, working together to provide a cohesive experience. Disentangling them is essential for understanding how they operate as drivers (Berkman et al., 2014; Chavez et al., 2024).

Facial processing guides our experience day-to-day interpersonal interactions and thus has been of special interest within social and cognitive neuroscience. The advent of functional magnetic resonance imaging (fMRI) allowed a shift from simply locating brain regions to assigning function. The changing blood oxygen levels became a secondary, but reliable predictor of brain activation (Ogawa et al., 1992). Using a statistical model, voxel changes in activation magnitude could be correlated to time points of interest both within and between subjects, and thus univariate analysis was born (Friston et al., 1994). The fusiform face area (FFA) was discovered to be the central facial processing region by Kanwisher, McDermott, and Chun (1997). Univariate analysis of fMRI confirmed this specialized network of nerves activate above baseline when a subject is presented with facial stimuli (Ogawa et al., 1992; Smith et al., 2004a).

An issue univariate fMRI analyses is that they focus on narrow regions showing changes in activation magnitude in highly localized regions that may not capture how the brain computes information across integrated brain regions. To address some of these limitation, Haxby and colleagues (2001) introduced the idea of using multivariate pattern analysis (MVPA) to analyze fMRI data. Rather than studying cognitive processes by changes in activation magnitude, MVPA seeks to uncover distributed patterns of activation across integrated areas. Haxby et al. (2001) found evidence of distributed and overlapping networks involved in both facial and object recognition in the ventral temporal cortex. Their results demonstrated how machine learning models accurately predicted the stimulus category based on neural activation patterns from both maximally and minimally responsive region. These findings had likely previously been masked by univariate analyses. MVPA sensitivity to distributed information found in our representational systems may be more appropriate for the naturalistic, highly variable stimuli inherent in social neuroscience (Fusar-Poli et al., 2009; Weaverdyck et al., 2020).

Multivariate analyses have since been enhanced to increase their usefulness for analysis of neural representational systems. Representational similarity analysis (RSA), developed by Kreigeskorte, Mur, and Bandettini (2008), quantifies the “distance” between representational patterns. The distance is a proxy for the degree of similarity. A considerable number of cognitive neuroscience experiments are aimed towards discovering how the brain reacts to various stimuli, designed with neural activation patterns as the dependent variable (Popal et al., 2019). Results typically show how activating distinct cognitive processes generate specific neural activation patterns, often within a localized region. Only recently has there been research to support the reverse inference that activation of these specific neural patterns can predict distinct cognitive processes (Chavez et al., 2024). Machine learning models are becoming increasingly vital in this area, utilizing neural activation patterns to quantify and predict the significance the similarities between experimental patterns and machine-generated model patterns.

The present study aims to expand on the understanding of what drives the neural representation of dynamic emotion perception using both naturalistic stimuli and the application of RSA on fMRI data. To address this question, we collected data from participants told to code the valence of the following set of stimuli depicting both positive and negative emotions. The recorded stimuli consisted of different volunteers making happy or angry faces directed toward or away from the camera. Many previous studies have explored dynamic facial processing with static, animated or computer-generated images. However, there is a lack research utilizing more ecologically relevant stimuli (i.e. real human faces). By incorporating real-life volunteers and RSA we aim to close the gap between experimental results and the reality of emotion perception. Our focus was on three possible aspects of these stimuli as potential drivers of neural representation: key person identifiers, emotional valence, and attentional orientation. We hypothesized the three factors would interact in distributed and overlapping neural representations.

## Methods

### Participants

Thirty-three subjects between the ages of 18 and 19 (22 female) were recruited from the local community at Dartmouth College in Hanover, New Hampshire. All subjects were screened to be right-handed and self-reported no current or history of psychiatric or neurological conditions. Subjects underwent an MRI protocol which included functional, diffusion weighted, high-resolution anatomical scans. Quality control procedures revealed that each subject in had head movement less than 3mm (corresponding to the voxel size) across all runs. Each anatomical scan was inspected visually to ensure there were no gross distortions, misalignments, or poor subcortical segmentation results. Thus, all 33 subjects were included in the final analysis. Subjects gave informed consent in accordance with the guidelines set by the Committee for the Protection of Human Subjects at Dartmouth College and received course credit or were paid for their participation.

### Image Acquisition

Magnetic resonance imaging was conducted with a Philips Achieva 3.0 Tesla scanner using a 32-channel phased array coil. Structural images were acquired using a T1-weighted MP-RAGE protocol (220 sagittal slices; TR: 8.176 ms; TE: 3.72 ms; flip angle: 8°; 1 mm isotropic voxels). Diffusion weighted images were collected using 70 contiguous 2-mm-thick axial slices with 32 diffusion directions (91ms TE, 8848 TR, 1000 s/mm2 b-value, 240mm FOV, 90° flip angle, 1.875 × 1.875 × 2 mm voxel size). Two diffusion scans were acquired per subject. Functional images were acquired using a T2*-weighted echoplanar sequence (TR: 2000 ms; TE: 35 ms; flip angle: 90°). For each participant, four runs of 168 whole-brain volumes (35 axial slices per whole-brain volume, 3×3×3.486 voxel size) were collected.

### Stimuli

Stimuli presented during the functional runs consisted of 2 second video clips of 16 volunteers making emotional expressions directed either towards or away from the camera. The video clips were recorded using a Canon VIXIA HF G30 HD Camcorder and edited using Apple iMovie software. The volunteers consisted of an ethnically diverse sample (9 White; 2 Asian; 2 Hispanic/Latino; 1 Black; 2 other) of equal number of males and females. Critical for this study was manipulation of self-engagement to the perception of each of the emotional facial expression. To accomplish this, volunteers were faced at a 45° angle pointing away from the camera at the beginning of each clip. For half of the clips, volunteers were instructed to make either a happy or angry facial expression while remaining facing at the angle directed away from the camera. During the other half of the clips, they were instructed to turn to the camera before making either a happy or angry facial expression. This combination results in a 2×2 design with emotion type and direction as each factor. Importantly, beginning each clip at the 45° angle ensured that information about the direction of the emotion could not be inferred from the angle of the shoulders at the beginning of the clip without seeing the volunteer in motion. A total of 16 volunteers, evenly split between males and females (9 White, 2 Asian, 2 Hispanic/Latino, 1 Black, and 2 other), created the stimulus videos. Figure 1 shows an example from one such volunteer. Each two-second video featured a single volunteer performing one of the following conditions: a happy face made while facing away from the participant, a happy face made while turning toward the participant, an angry face made while facing away from the participant, and an angry face made while turning toward the participant. All videos were filmed with volunteers starting 45° away from the camera, so emotional valence couldn’t be inferred from body placement alone. This resulted in four conditions for each of the 16 targets, yielding 64 naturalistic video stimuli. Researchers have found that naturalistic stimuli provide more robust signaling information, capturing more facial processing regions of interest than static stimuli (Fox et al., 2009). Using naturalistic stimuli rather than static images or drawings, as some other research has done, also yields more ecologically relevant results (Berkman et al., 2014).

**Figure 1.**
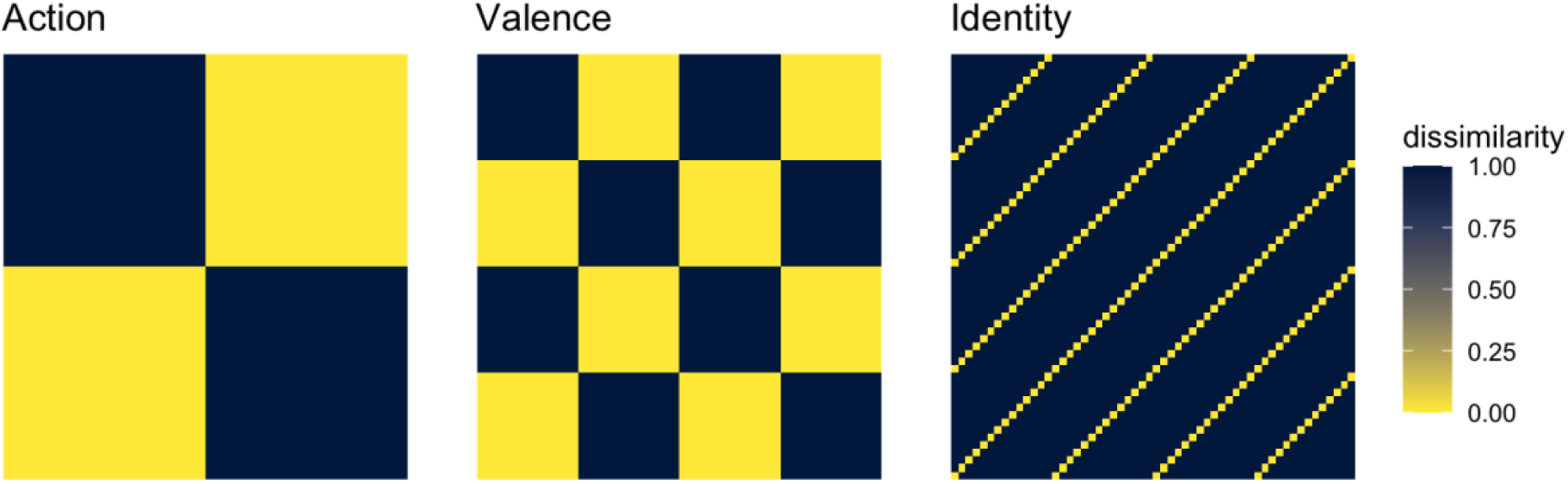
Hypothesized RDMs representing distinct hypotheses for neural encoding of action, valence, and identity. The action model predicts neural similarity based solely on the direction of gaze (toward self vs. away from self). The valence model predicts that stimuli with similar emotional affect (positive vs. negative) will elicit similar neural patterns, regardless of other conditions. Finally, the identity model predicts neural similarity based on target identity alone. Dissimilarity is represented on a scale from 0 (yellow: identical patterns) to 1 (navy: maximally dissimilar patterns). These models serve as the computational framework against which recorded neural data is compared using RSA.

### Procedure

During the fMRI scan, subjects were shown the video clips on a screen adjacent to the rear scanner bore through a mirror attached to the head-coil. Each video clip was presented for a single 2000 ms TR in a rapid event-related design. Inter-trial intervals consisting of a fixation cross for 2000 ms were pseudo-randomly interspersed to introduce jitter into the fMRI time series. Each stimulus category stimulus identity was pseudo-randomly presented once in each of the four runs. Subjects were told they would be viewing a series of people making facial expression from different angles. During the presentation of the clips, subjects responded via button-box as to whether the emotion displayed in the clip was positive or negative. Having subjects explicitly focus on the emotion presented in the clips encouraged them to process the emotional content of the faces while also being naïve to the purpose of the direct vs. indirect views. Trials were then sorted into the four task conditions and were then used as regressors in the fMRI time-series.

### Functional magnetic resonance imaging preprocessing

The fMRI data were analyzed using FSL (Smith et al., 2004b). Preprocessing of the fMRI data followed a standard procedure. First, all slices were interpolated to a common time point (i.e., slice-time correction) to correct for differences in slice acquisition. Next, images were smoothed using a Gaussian kernel of 4 mm FWHM, mean-based intensity normalization of all volumes by the same factor, and high-pass temporal filtering (Gaussian-weighted least-squares straight line fitting, with sigma = 90.0 s). Time-series statistical analyses were carried out using local autocorrelation correction. A two-step normalization process was performed using linear registrations with FLIRT by aligning functional data to each subject’s anatomical scan before registering it to the MNI template using non-linear registration with FNIRT. The general linear model was then employed to measure activation patterns to each trial per run individually using a least-squares-single modeling procedure (Mumford et al., 2014). Normalized (i.e., z-scored) voxel responses for each condition were extracted from each parcel of the 400 regions in the Schaefer parcellation atlas (Schaefer et al., 2018). Consistent with previous studies using fMRI to estimate multivariate similarity patterns (Kriegeskorte et al., 2008), we used Spearman rank correlation distance (e.g., 1 – Pearson-*R*) to compute the dissimilarity between neural response patterns among each of the conditions (i.e., the self and each of the other people in the group) using tools from the nilearn package in Python (Abraham et al., 2014). These values were used as the estimates of neural similarity in each brain parcel that were then compared against the hypothesized action, valence and identity representational dissimilarity matrix (RDM) models (Figure 2).

**Figure 2.**
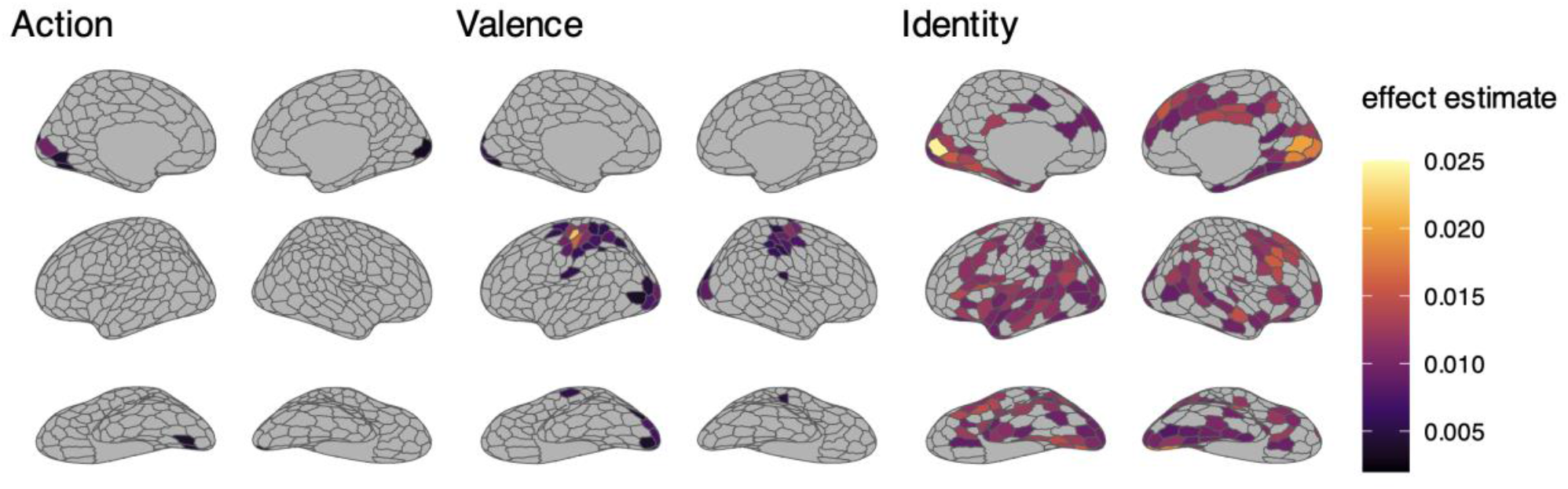
Cortical mapping of the result of the representational similarity analysis of action, valence, and identity models. The action map illustrates regions where the neural pattern similarity is significantly driven by gaze direction. The valence map illustrates regions that are significantly predicted by the emotional affect of the stimuli. The identity map highlights cortical areas where representational patterns differentiate between the identities of individual stimuli.

Action, valence, and identity RDM models were then related to neural-similarity measures at every parcel within each participant using a representational similarity analysis (Kriegeskorte et al., 2008). We fitted a nested multilevel model using all three off-diagonal RDMs as predictors of brain-response similarity within random intercepts for runs nested within subjects using the lme4 package in R (Bates et al., 2015). The significance of each RDM was calculated using Satterthwaite estimated p values for every parcel (Kuznetsova et al., 2017). Finally, whole-brain significance tests were corrected for multiple comparisons using a Bonferroni correction across parcels for each.

## Results

MVPA of fMRI data was conducted independently across 414 parcels. The goal of the RSA was to determine where patterns of activity were most significantly alike the RDM models generated for each condition (action, valence, target identity), while controlling for the other two. Thus, revealing where each condition acts as a driver for our neural representation of facial perception. The results of the analysis are shown in Figure 2. The resulting patterns are minimally overlapping, with the most widespread effects occurring within the identity category. Tables 1-3 show a complete list of the significant parcels. Importantly, these results do not tell us whether a region is more or less active. Rather, the data show the regions that encode necessary information related to each condition, which is integrated with information from other regions that contribute to our ability to process facial information.

**Table 1.**
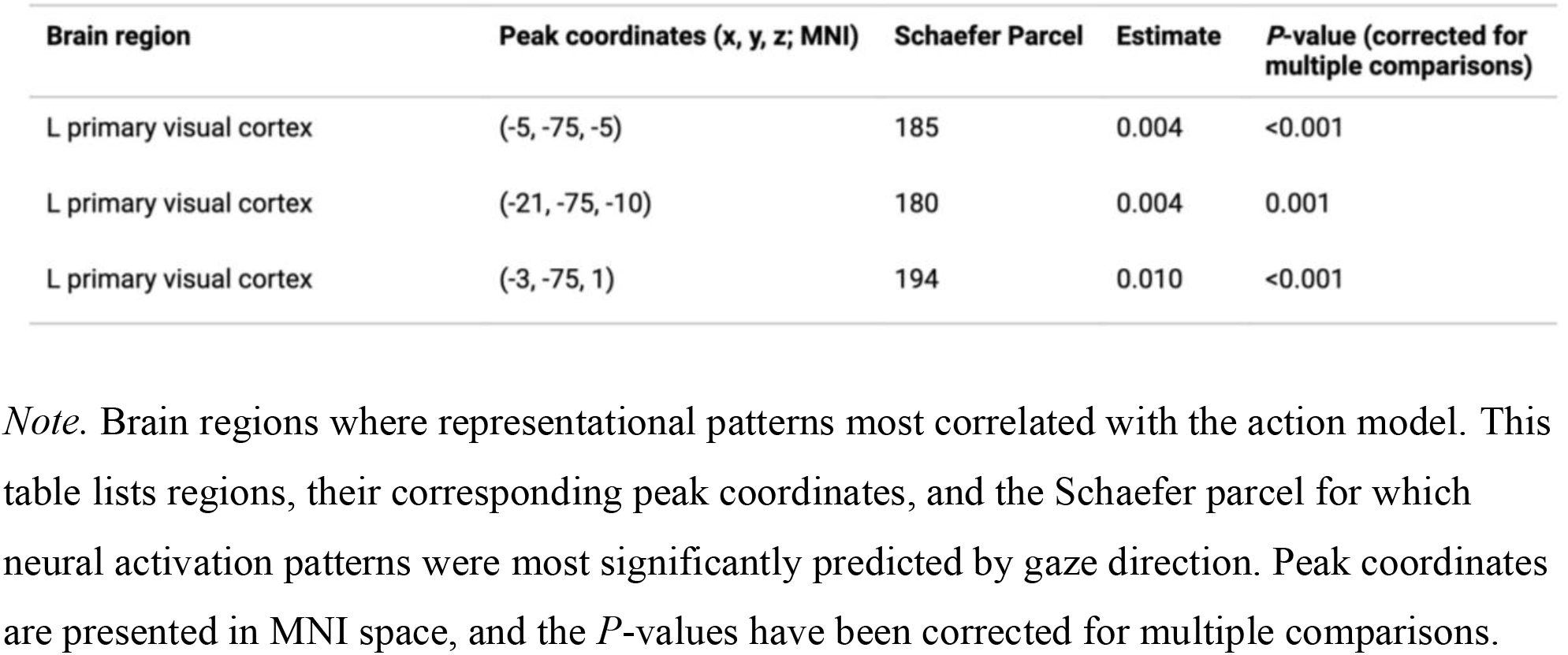

**Table 2.**

| Brain region | Peak coordinates (x, y, z; MNI) | Schaefer Parcel | Estimate | P-value (corrected for multiple comparisons) |
| --- | --- | --- | --- | --- |
| L somatosensory network | (-28, -17, 71) | 149 | 0.012 | <0.001 |
| R somatosensory network | (28, -11, 71) | 329 | 0.010 | <0.001 |

*Note.* Brain regions where representational patterns most correlated with the valence model. This table lists regions, their corresponding peak coordinates, and the Schaefer parcel for which neural activation patterns were most significantly predicted by emotional affect. Peak coordinates are presented in MNI space, and the *P*-values have been corrected for multiple comparisons.

**Table 3.**

| Brain region | Peak coordinates (x, y, z; MNI) | Schaefer Parcel | Estimate | *P*-value (corrected for multiple comparisons) |
| --- | --- | --- | --- | --- |
| L dorsal medial prefrontal cortex | (-3, 48, 36) | 23 | 0.010 | 0.015 |
| R dorsal medial prefrontal cortex | (6, 48, 36) | 269 | 0.015 | <0.001 |
| L posterior cingulate cortex | (-3, -39, 25) | 76 | 0.013 | 0.015 |
| R posterior cingulate cortex | (3, -33, 26) | 271 | 0.013 | 0.001 |
| L temporoparietal junction | (-50, -51, 13) | 108 | 0.011 | 0.005 |
| R temporoparietal junction | (50, -44, 16) | 344 | 0.010 | 0.008 |
| L amygdala | (-25, -11, -21) | 405 | 0.010 | <0.001 |
| R amygdala | (25, -11, -21) | 412 | 0.009 | 0.001 |
| L fusiform face area | (-40, -61, 13) | 138 | 0.010 | 0.018 |
| R fusiform face area | (40, -61, 13) | 375 | 0.009 | <0.001 |

*Note.* Brain regions where representational patterns most correlated with the identity model. This table lists regions, their corresponding peak coordinates, and the Schaefer parcel for which neural activation patterns were most significantly predicted by target identity. Peak coordinates are presented in MNI space, and the *P*-values have been corrected for multiple comparisons.

### Action

RSA revealed several parcels of interest that store our neural representation of action (ie. look vs away). All parcels fell within the primary visual cortex (V1) of the occipital lobe, which is associated with early, simple visual processing. A summary of such brain regions is provided in Table 1.

### Valence

Results in the valence condition partially overlap the look category. However, several parcels extend beyond early V1 and begin to enter more specialized visual system regions (Fig. 4). We also observe highly statistically significant input from both the right and left somatosensory networks (*b*=0.012, *P* <0.001; *b*=0.010, *P* <0.001). The primary task required participants to code the stimuli valence, thereby contributing to this effect. Notably, we do not see a significant contribution from the limbic system, the brain’s center for emotional processing.

### Identity

The identity condition resulted in the most widely distributed representation of facial processing. Results included several parcels that overlapped both the look and valence conditions. Consistent with prior research, the activation patterns within the fusiform face area are predictive of facial identity (*b*=0.009, *P*=<0.001). Most interesting is the number of parcels within the default mode network (DMN), a network of brain regions that is consistently suppressed during goal-directed tasks. These regions include the right dorsal medial prefrontal cortex (R dmPFC) and R posterior cingulate cortex (R PCC) (*b*=0.015, *P*=<0.001; *b*=0.013, *P*=0.001). A summary of select DMN and supporting regions is in Table 3.

## Discussion

Effective social interaction requires the use of overlapping, widely distributed representational systems, several of which contribute to our dynamic facial processing ability. This research used naturalistic stimuli to determine which aspects of stimuli drive our encoded neural patterns. We focused on three possible conditions as drivers: action (as a representation of social attention), valence, and stimulus facial identity. Using RSA of fMRI data, we compared hypothetical models of neural patterns with real, experimentally derived patterns to begin disentangling the conditions. Analysis showed that each of the three proposed conditions plays a role in driving our neural representations in more than one brain parcel. Thus, supporting the conjecture that the brain is not a one-to-one processing unit, but rather stores information in diffuse patterns. Results show that facial identity, as a driver, is encoded in the most widely distributed pattern across all conditions. Others have also found similar results within the ventral stream, anterior temporal lobe, and anterior fusiform gyrus (Goesaert & Op De Beeck, 2013; Nestor et al., 2011). Regions identified in this experimental context (Fig. 4) reveal that identity is something the brain organizes more globally.

Areas of significance within the active condition included V1, the early visual cortex responsible for lower-level visual processing. An initial interpretation of this data is that stimuli that are actively turning appear more similar, not because of the overall perceived image, but because of a simple identifier, such as the observed lines or sharp corners. All stimuli were recorded at 45°, so placement alone did not indicate whether the stimuli would face away from or turn toward the camera. The early visual cortex is one such area where there is overlap between the action and identity conditions, indicating that they act independently as significant drivers of facial perception. One subset of facial perception is facial recognition, the ability to differentiate between two distinct faces. The consensus has long been that holistic processing, or processing of the face as a single, integrated whole, is an accurate predictor of facial recognition. Evidence now suggests that feature-based processing may be a more accurate predictor of early facial recognition (Goesaert & Op De Beeck, 2013). New findings, together with our results, may explain why action as a driver of facial processing appears confined to these early visual areas.

Results revealed two main regions in which valence drove our neural representation: V1 and the primary sensorimotor cortex (SM1), a broad area spanning the central sulcus responsible for both sensory and motor functions. The parcels within V1 were slightly past the early visual cortex, now entering ventral temporal areas and partially overlapping the fusiform gyrus. As discussed, this brain region plays a central role in encoding our neural representation of faces (Fusar-Poli et al., 2009; Ganel et al., 2005; Kanwisher et al., 1997). However, because RSA captures multidimensional neural patterns, our results indicate the FFA is not the only region contributing to any of the three conditions. Valence also drove neural representation within SM1, likely due to the scanner task. Participants were instructed to code each stimulus for valence using a button placed in their right hand, thereby reinforcing neural dissimilarities between happy and sad faces. This likely explains why we see significant results within SM1, a confound of the experimental design.

It is surprising to see a lack of subcortical representation from regions within the limbic system. This may be because some structures, such as the amygdala, habituate quickly to emotion-based tasks (Breiter et al., 1996). Suggesting that there was no significant change in neural patterns between happy and sad stimuli. Without change, happy stimuli will look the same as sad stimuli, at least within the amygdala, which is incompatible with the hypothetical model generated.

As mentioned, the most numerous and distributed effects were observed in the target condition. It is clear from these results that more areas of the brain are involved in our neural representation of facial perception, particularly when considering identity alone. Unsurprisingly, there is some overlap with the fusiform face area. However, there are also several midline cortical regions, such as the medial prefrontal cortex (mPFC), posterior cingulate cortex (PCC), and the temporoparietal junction (TPJ). These regions are included in the DMN. Originally discovered due to its inactivity during goal-directed tasks (Raichle et al., 2001), further research has confirmed that the default mode is not simply our “baseline.” Rather, these brain regions are imperative for contributing to our understanding of self *and* others (Schilbach et al., 2008). Essentially, when our brains are not actively processing external stimuli, these regions enter a self-referential, daydreaming state.

Several significant parcels overlap the mPFC, and recent research shows this region acts as a hub of the DMN (Li et al., 2014). The ventral mPFC connects to emotional processing regions, such as the amygdala, working with the PCC to formulate our empathy response. Meanwhile, the anterior and dorsal mPFC primarily help us delineate self from others (Li et al., 2014). The mPFC and TPJ appear to be the primary regions involved in global switching between social and non-social cognitive processes, and more specific theory of mind tasks— though their exact roles are the subject of much debate among social neuroscientists. Theory of mind is our ability to interpret another person’s cognitive state based just on their behavior, or facial expression, in this case. It is evident from the many cognitive processes contained within the DMN that target identity is an incredibly complex condition. The hypothetical model can only consider stimulus identity at face value. The brain may be encoding more, ascribing mental states, and drawing connections between self and other.

### Future Directions

There were several limitations of this experiment due to my use of secondary data. In the future, I might alter the experimental design to more directly target the three proposed conditions (action, valence, and identity). For example, including stimuli designed to elicit a stronger, more varied emotional response from the participant may elicit activity in subcortical and limbic areas. I may also include more sub-conditions within identity that factor in stimulus emotion and action to better tease which aspects of identity drive which areas of the DMN. Finally, one could switch the participant task from categorizing valence to categorizing turning action (none or toward self) to ensure we see a similar pattern of activation as resulted in the present design.

## Conclusion

In conclusion, the results of this analysis reveal that determining who you are looking at requires more brain areas than determining the stimulus’s emotional valence or social orientation. Our neural representations of facial perception in DMN regions are driven by stimulus identity. In other words, when your eyes catch another’s in the grocery store, the activation patterns across your brain store information regarding who that person is in relation to you, considering whether you’ve seen them before, and ascribing cognitive processes based on their expression. This finding may shape how researchers design and analyze future social perception tasks. If one doesn’t adequately account for facial identity you may miss some of what is driving the main effect of the experiment or conflate results.

